# VEGF-A, PDGF-BB and HB-EGF engineered for promiscuous super affinity to the extracellular matrix improve wound healing in a model of type 1 diabetes

**DOI:** 10.1101/2021.06.21.449313

**Authors:** Michael JV White, Priscilla S Briquez, David AV White, Jeffrey A Hubbell

## Abstract

Chronic non-healing wounds, frequently caused by diabetes, lead to lower quality of life, infection, and amputation. These wounds have limited treatment options. We have previously engineered growth factors to bind to exposed extracellular matrix (ECM) in the wound environment using the heparin-binding domain of placental growth factor-2 (PlGF-2_123-144_), which binds promiscuously to ECM proteins. Here, in the type 1 diabetic (T1D) NOD mouse model, engineered growth factors improved both re-epithelialization and granulation tissue formation. Engineered growth factors were even more potent in combination, and the “triple therapy” of vascular endothelial growth factor-A (VEGF-PlGF-2_123-144_), platelet-derived growth factor-BB (PDGF-BB-PlGF-2_123-144_), and heparin-binding epidermal-growth factor (EGF-PlGF-2_123-144_) both improved wound healing and remained at the site of administration for significantly longer than wild-type growth factors. In addition, we also found that changes in the cellular milieu of a wound, including changing amounts of M1 macrophages, M2 macrophages and effector T cells, are most predictive of wound healing success in the NOD mouse model. These results suggest that the triple therapy of VEGF-PlGF-2_123-144_, PDGF-BB-PlGF-2_123-144_, and EGF-PlGF-2_123-144_ may be an effective therapy for chronic non-healing wounds in that occur as a complication of diabetes.

## Introduction

Diabetes is a major health scourge that affects more than 415 million people worldwide. One in eleven adults has diabetes, and 12% of all global health expenditures are related to diabetes [1], including chronic non-healing diabetic wounds. The risk for diabetic patients to develop lower extremity non-healing wounds ranges from 15 to 25% during their lifetime [2, 3], and about 33% of the direct costs of diabetes are linked with the treatment of diabetic foot ulcers [4, 5].

Type 1 diabetes (T1D) affects 1.25 million Americans, 5% of diabetics overall, and primarily manifests in children [6]. T1D is the most prevalent form of diabetes in youth and accounts for more than 85% of young diabetic patients [6]. Further, T1D has been increasing in global incidence by 3% annually [6]. Although T1D has only 5-10% of the prevalence of type 2 diabetes (T2D), the health complications for T1D are more severe [7, 8], with persons with T1D heal both acute and chronic wounds poorly, independent of their glycemic control [9].

Chronic non-healing wounds affect more than 6.5 million US patients per year, which is approximately 2% of the population, and cost more than $ 25 billion to treat [10]. Compared to healthy wounds, chronic non-healing wounds exhibit poor resolution of the inflammatory phase of wound healing [11]. This includes impaired leucocyte recruitment and phagocytic activity and increased concentrations of pro-inflammatory cytokines [12, 13].

The extracellular matrix (ECM) is a highly dynamic structure that modulates cell proliferation, migration, and differentiation during the course of wound healing [14]. During wound healing, immune cells and fibroblasts secrete various growth factors (GFs) in the wound, most of which interact with ECM before finding their cognate cell surface receptors. The ECM sequesters GFs, which creates a slow-release reservoir of GF for local signaling [15, 16]. ECM composition and secreted GFs create a signaling microenvironment that tightly controls cellular responses in the healing wound [14]. This signaling environment changes as the wound heals, as do the types of cells interacting in the wound [17].

Due to an excessively proteolytic environment, chronic non-healing wounds have a fragmented ECM structure, which results in poor sequestration of GFs. In addition, the high protease activity in chronic wounds digests GFs and GF receptors at an increased rate [18]. Together, the dysregulated ECM and increased proteolysis of chronic non-healing wounds disorganize cellular behavior (i.e., migration, proliferation, differentiation) and prevent wound resolution, since proper cellular orchestration is key to resolving each stage of wound healing [14].

GFs are potent biological signals that can instruct cell morphology, metabolism, and differentiation. GFs have been hindered as medical treatments due to off-target signaling [14, 19], low signaling efficacy, and high expenses due to the high concentration of GFs necessary to overcome the off-target signaling and low signaling efficacy [20, 21].

Platelet-derived growth factor-BB (PDGF-BB) is one of the first GFs to be released into the wound environment by platelet degranulation and is subsequently secreted by monocyte-derived cells, fibroblasts, and endothelial cells in later stages of wound healing [22-24]. PDGF-BB thus plays an important role in multiple subsequent stages of wound healing. Particularly, PDGF-BB promotes the development of granulation tissue within the wound via its effects on fibroblast proliferation, mesenchymal stem cell (MSC) recruitment and ECM production [22].

Vascular endothelial growth factor-A (VEGF-A) is responsible for angiogenesis in wounds by triggering the angiogenic cascade [25, 26]. On a cellular level, VEGF interacts with monocyte-derived cells and endothelial cells through the VEGF-receptor-2 (VEGFR2) to create new blood vessels within granulation tissue [25, 27]. Like PDGF-BB, VEGF-A is released through subsequent wound stages by multiple cell types, including platelets, monocyte-derived cells, fibroblasts and endothelial cells [27].

Heparin-binding epidermal-growth factor (HB-EGF) has increased secretion in damaged skin [28] and is secreted by macrophages [29]. HB-EGF operates through epidermal growth factor receptor(EGFR, also called human epidermal growth factor receptor HER1) [28], affects keratinocyte proliferation and migration (dependent on dose and means of presentation [30-32]), and accelerates wound re-epithelization [23]..

While several GFs play roles in wound healing [22, 33], PDGF-BB, VEGF-A, HB-EGF are especially promising as drug targets due to their signaling potency and ability to interact with multiple cell types in the wound environment [23, 34].

Recombinant GFs have been explored in human trials as topical treatments for diabetic foot ulcers: PDGF-BB [35], VEGF-A (NCT00351767), and HB-EGF (NCT01629199). Of these, only PDGF-BB has thus far been approved for human use as the topical gel Regranex. However, Regranex, which is used at a rather high dose of 7 µg/cm^2^ and is applied daily for up to 2 weeks [36], carried a warning about potential carcinogenic effects due to the potential for recombinant PDGF-BB to escape into the bloodstream and cause distant tumors through off-target signaling [37].

Thus, the development of a delivery approach that reduces off-target signaling of GFs and increases the local concentration of GF at appropriate sites of tissue regeneration could be highly beneficial for GF-based tissue regeneration therapies. Previously, we engineered growth factors (eGFs) that promiscuously bind to ECM proteins in wounds [20], by fusing them to the heparin-binding domain from placental growth factor-2 (-PlGF-2_123-144_). These eGFs speed wound healing, can be used at lower concentrations, and thus may be safer than wild-type (WT) GFs [20]. Indeed, we showed that the eGFs have increased retention at the site of application as a result of binding to the ECM [20]. Increased retention at the site of application reduces the risk of eGFs escaping into the blood or lymphatic system and causing off-target effects.

Wound healing is a complex process in which a multitude of cell types migrate, differentiate, and proliferate in an environment of complex secreted signals and ECM surfaces. Further, this cellular milieu changes as the wound heals, with each successive change bringing the wound closer to resolution through the inflammatory, proliferative, and remodeling phases of wound healing [38, 39]. Thus, understanding the cellular milieu of wounds is essential for a mechanistic understanding of eGF’s effect on wounds.

The initial inflammatory stage of wound healing are characterized by neutrophil infiltration, and neutrophil persistence in wounds is key to the inflammation within the wound. This stage of wound healing lasts for a period of hours up to days after wounding [17].

The proliferation phase of wound healing begins after the inflammatory stage begins, and is characterized by infiltration of mesenchymal stem cells, smooth muscle cells, endothelial cells, and macrophages [20]. The tissue remodeling phase usually begins 1-3 days after wounding [40], and includes the T cell response to the wound [17]. Macrophages are key effectors of wound healing [41], secreting several cytokines and GFs, and mediating the differentiation and proliferation of several types of cells in the wound.

Macrophages in a wound differentiate into several different types: classically-activated macrophages (M1 macrophages) are key to perpetuating inflammation and phagocytosing cell debris in the wound [42]. Alternatively-activated (M2) macrophages are responsible for tissue remodeling [42]. Arginase is an enzyme that catalyzes the production of ornithine and urea, and arginase-positive macrophages promote wound healing [43]. The presence and appropriate differentiation of macrophages within a wound is essential for the restoration of tissues, and arginase-positive macrophages are necessary for proper wound healing [43]. Underscoring macrophage importance in healing wounds, depletion of macrophages from a wound inhibits wound healing [44].

Non-hematopoietic cells are also key to wound regeneration. Particularly, endothelial cells are key to lining new blood vessels, and mesenchymal stem cells form the basis of much of wound healing through their ability to differentiate into multiple cell types [17].

If wounds remain unhealed for enough time, the adaptive immune system can begin to affect the outcome of the wound [17]. Amphiregulin is an autocrine GF, and amphiregulin-positive tissue-repair T cells are a newly discovered class of T cells that are important for wound healing [43, 45]. T-regulatory cells (Tregs) are anti-inflammatory cells, and removal of Tregs slows wound closure [46].

The diabetic-specific problems of non-healing chronic wounds (abnormal glycation in the wound environment, increased proteolytic activity, and poor ECM sequestration) considerably reduce the signaling efficacy of locally produced GFs on the cell behavior in a wound (proliferation, migration, and differentiation). This cellular dysregulation disrupts the resolution of the inflammatory, proliferation, and remodeling phases of wound healing [38, 39]. Because eGFs bind strongly to exposed ECM, they are sequestered in the ECM of chronic non-healing wounds and protected from proteolysis better than are recombinant WT GFs. This dysregulated environment makes eGFs a promising treatment for delivering GFs potent signaling to the wound environment.

## Results

We previously observed that the PlGF-2_123-144_ domain was a promiscuous binder of multiple ECM proteins [20]. Co-delivery of engineered VEGF-A-PlGF-2_123-144_ and PDGF-BB-PlGF-2_123-144_ improved re-epithelialization more than co-delivery of VEGF-A and PDGF-BB in the T2D db/db mouse model. Here, we sought to extend this work to a T1D model using the NOD mouse, and to explore addition of a third eGF, namely HB-EGF-PlGF-2_123-144_.

To model type 1 diabetic wound healing in manner relevant to a clinical setting, we allowed NOD mice to become diabetic spontaneously, then treated them until their glucose was sub-optimally controlled with insulin, then wounded them. To determine if GF or eGF (GF-PlGF-2_123-144_) can improve wound healing in the T1D NOD model, we added 200 ng of VEGF-A, PDGF-BB, and HB-EGF (GF and eGF variants) to wounded skin.

While no GF or eGF increased granulation tissue until 7 days (Figure 1A and C), co-administration of VEGF and PDGF-BB improved the re-epithelialization of NOD wounds at both 3 and 7 days (Figure 1B and D). The addition of HB-EGF resulted in an increase in granulation tissue as compared to wounds treated with fibrin alone after 1 week (Figure 1C). Treatment with the triple therapy (VEGF-A-PlGF-2_123-144_, PDGF-BB-PlGF-2_123-144_, and HB-EGF-PlGF-2_123-144_) increased both wound closure and granulation tissue compared to the NOD control mouse, but also compared to the non-diabetic NOD variant NOR mouse (Figure 1C and D). Further, the triple therapy eGFs outperformed the WT variants (Figure 1C and D) in both granulation tissue formation and wound closure. eGF triple therapy did not outperform GF therapy based on specific activity, because both GF and eGF both activate their receptors to equivalent extents (Figure 2A). eGFs are, however, retained in wounded tissue at higher concentrations for significantly longer time than the corresponding GF (Figure 2B). By way of comparison with our previous study [20] with VEGF-A-PlGF-2_123-144_, PDGF-BB-PlGF-2_123-144_ combination in the db/db T2D model, the eGF triple therapy led to more granulation tissue (Figure 1C) and more extensive would closure (Figure 1D) at 7 days than did the dual combination therapy. The eGF triple therapy was also more efficacious than HB-EGF-PlGF-2_123-144_ by this measure (Figure 1D). Figure 1E shows representative histology staining of wound sections.

**Figure 1:**
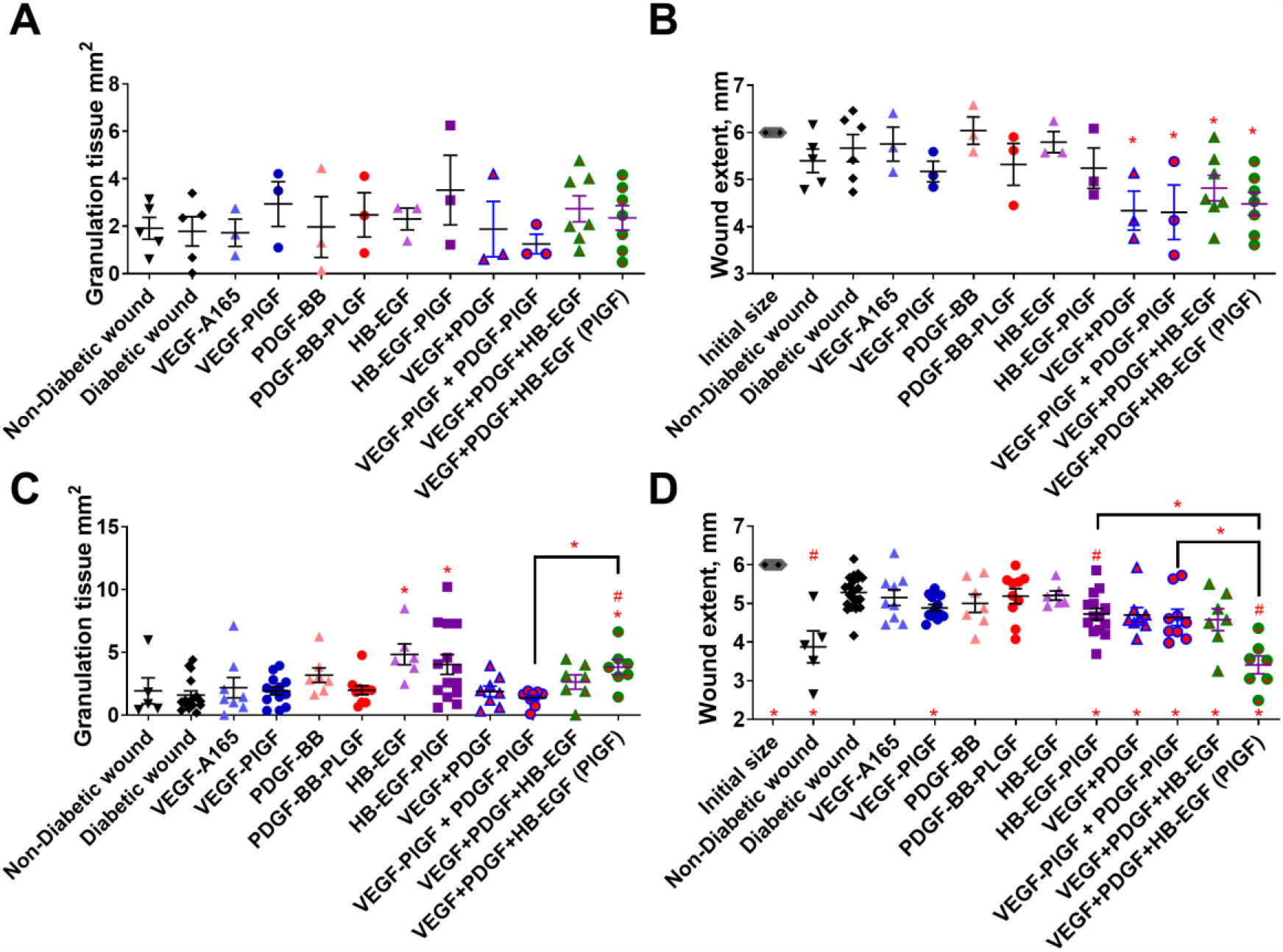
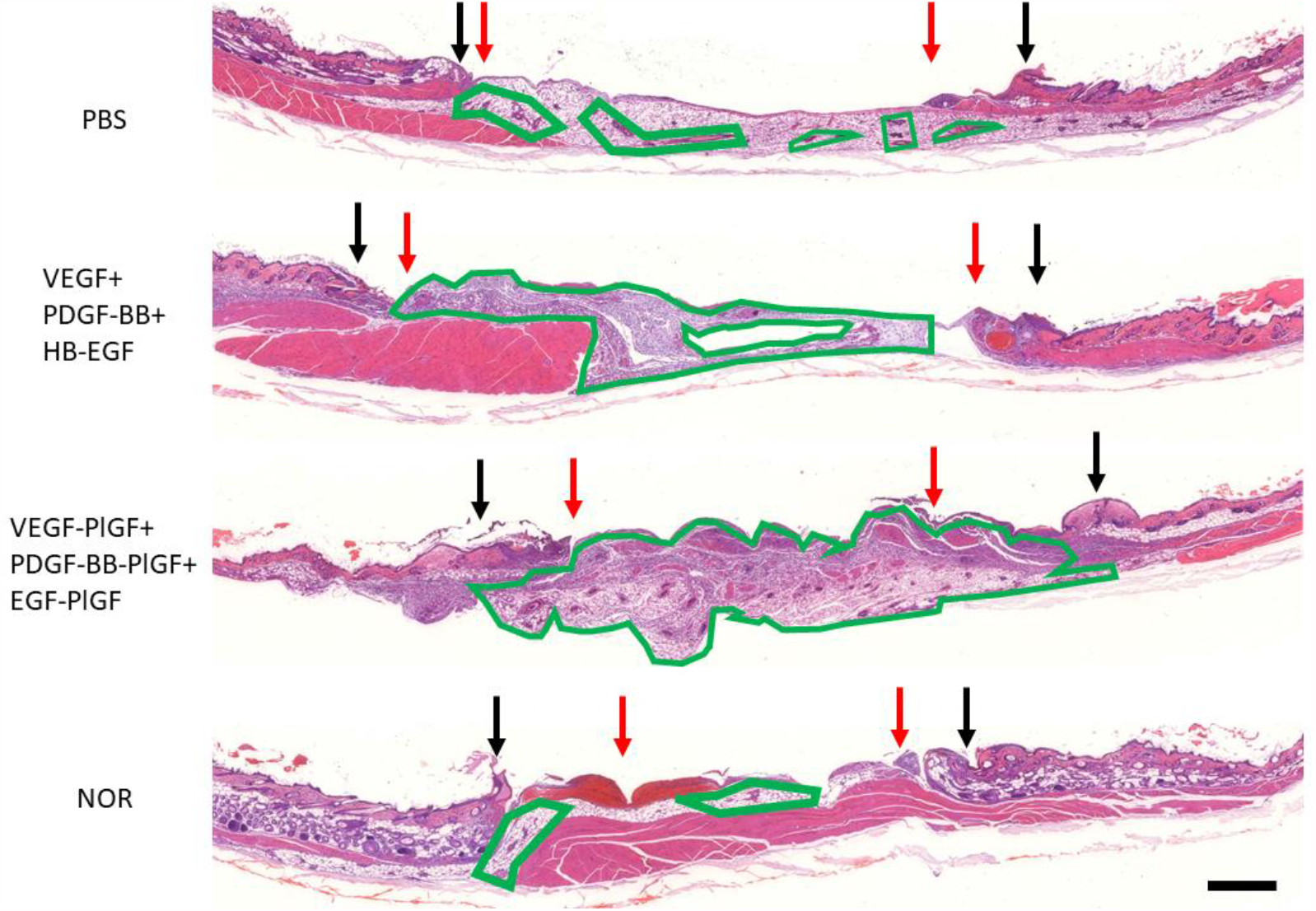
The triple therapy of VEGF-PlGF-2_123-144_, PDGF-BB-PlGF-2_123-144_, and HB-EGF-PlGF-2_123-144_ improves wound healing in the T1D NOD mouse model, as measured by increased granulation-tissue formation (A,C) and wound extent (diameter of unhealed wound normalized to the known length of the resection biopsy punch (12 mm) to account for folding or contracture of wound tissue, B, D). Timepoints assayed are 3 days (A,B) and 7 days (C,D). For wound extent, smaller numbers indicate more healing. (E) Representative images of healed wounds from 7 days, stained with hematoxylin and eosin. Black arrows indicate initial wound extent, red arrows indicate healed extent. Scale bar is 1 mm. n ranges from 3 to 20. * denotes comparison to the diabetic wound. # denotes a comparison between untreated NOD wound and untreated NOR wound, and the WT GF(s) are compared to their counterpart -PLGF-2_123-144_ variant(s), e.g., VEGF vs VEGF-PlGF-2_123-144_. * = P < 0.05, P < 0.01, P < 0.001, ANOVA + Student’s t-test for post-hoc for wound extent, Kruskal-Wallis + Mann-Whitney post-hoc test for granulation tissue. # = P < 0.05, P < 0.01, P < 0.001, Mann-Whitney for granulation tissue, Student’s t-test for wound extent. Student’s t-test for comparison between VEGF-PlGF+PDGF-PlGF and VEGF-PlGF + PDGF-PlGF + HB-EGF-PlGF.

**Figure 2:**
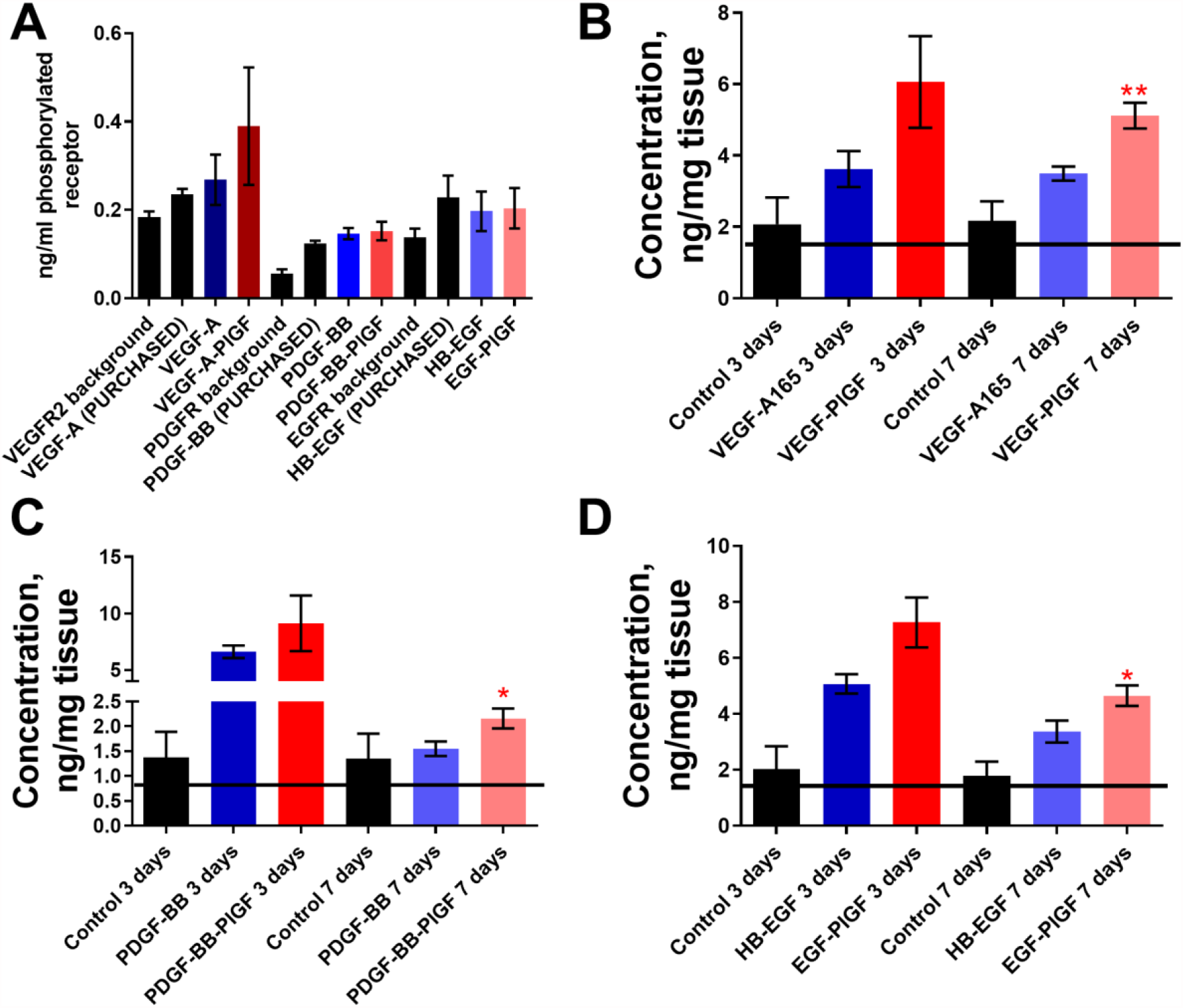
Growth factors fused with the -PLGF-2_123-144_ activate their cognate receptors at levels comparable to WT GF and remain in wound tissue longer than their wild-type (WT) counterpart. (A) 100 ng of VEGF-A-PlGF-2_123-144_, PDGF-BB-PlGF-2_123-144_ and HB-EGF-PlGF-2_123-144_ caused corresponding receptor phosphorylation (VEGFR2, PDGFR, EGFR respectively) at the indicated amounts over 10 minutes on HUVEC cells (for VEGF-A165) or fibroblasts (for PDGF-BB and HB-EGF). C) Concentrations in wounded skin tissue (ng/mg) for A) VEGF-A-PlGF-2_123-144_, D) PDGF-BB-PlGF-2_123-144_, and E) HB-EGF-PlGF-2_123-144_ after the indicated time points. n =7, * = P < 0.05, Mann-Whitney.

Interestingly, mouse-specific diabetic covariates (age, previous glucose high, glucose reading at the time of surgery, Table 1), are not predictive of wound healing outcomes (wound extent or granulation tissue), as indicated by a higher R^2^ value or by a lower p-value. Rather treatment with the triple therapy of eGFs (VEGF-A-PlGF-2_123-144_, PDGF-BB-PlGF-2_123-144_, and EGF-PlGF-2_123-144_) is more predictive of wound healing outcomes (Table 1). This suggests that the combination of these eGFs may be more useful treatments than GF in a clinical setting.

**Table 1:**
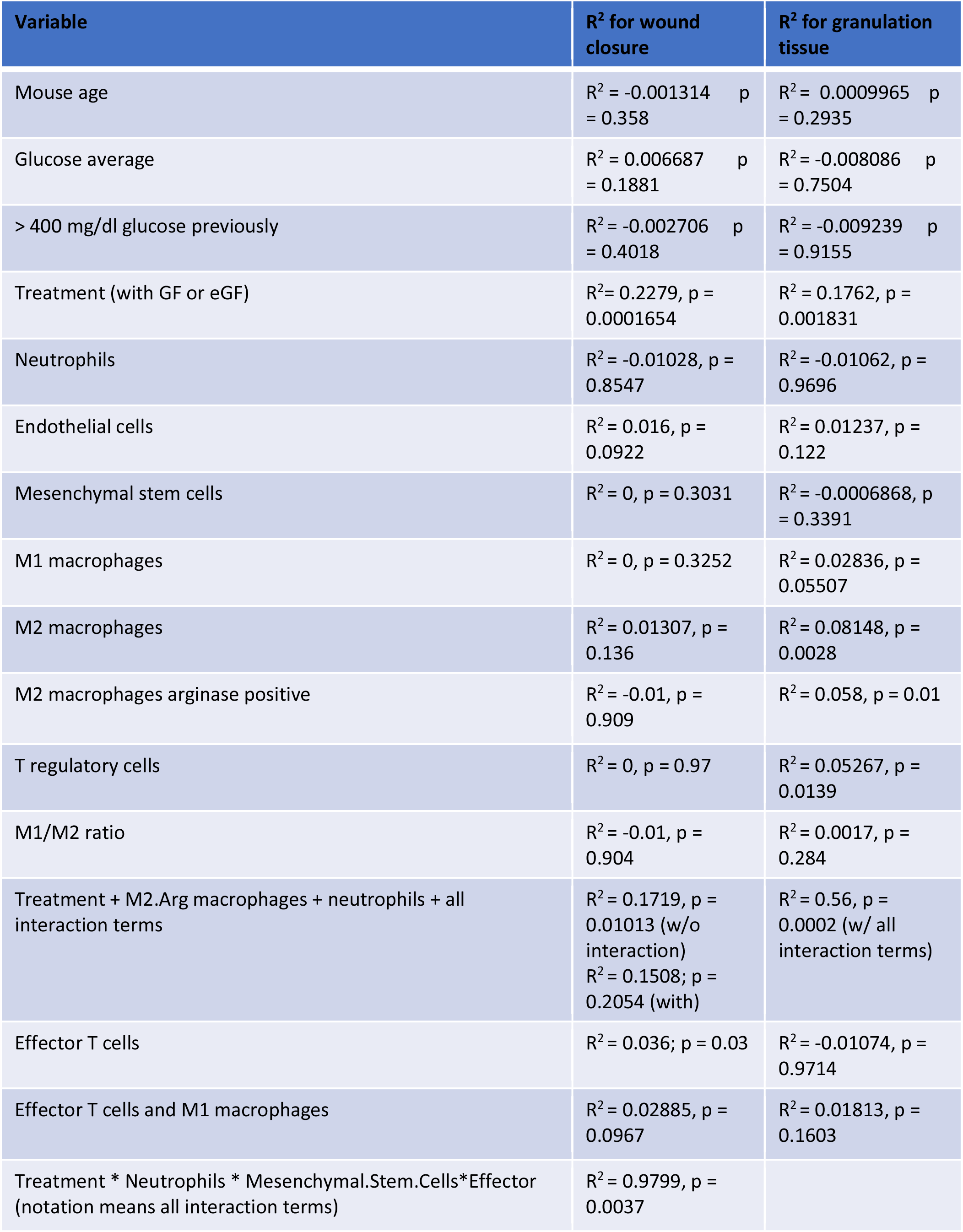
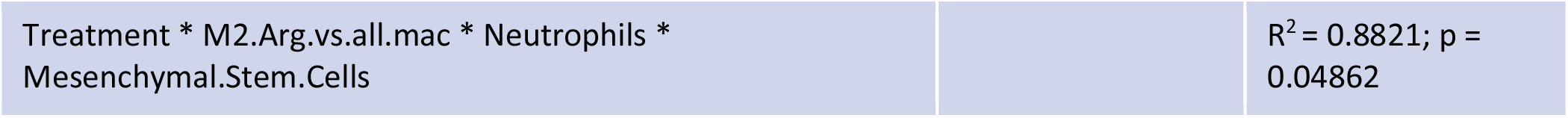
Statistical comparison of cell number in wound, correlated with wound healing outcome. Linear models using data on the cellular makeup of wounds (Figures 3-8) and mouse-specific covariates (age, average glucose at time of wounding, and previous high glucose reading >400 mg/dl) as explanatory variables and wound outcomes (wound extent or granulation tissue) as the response, using scatterplots to confirm the linear relationship. R^2^ measures the percentage of variance in the response variable explained by the explanatory variables. Adjusted R^2^ includes a penalty term for models with too many variables. * = P < 0.05, * * = P < 0.01, * * * = P < 0.001.

The cellular milieu of wounds changes through the stages of wound healing [17]. Non-healing diabetic wounds can become stuck in a non-resolving inflammatory cycle [17]. To determine if eGF treatment works through changing the cellular composition of the wounds, we analyzed cellular compositions of the wounds by flow cytometry at day 3 and 7 post-wounding. The triple therapy treatment significantly decreased the number of neutrophils at both time points, to levels comparable to non-diabetic NOR mouse wounds (Figure 3A and B). The eGFs of the triple therapy also outperformed the WT variant of triple therapy in neutrophil reduction (Figure 3A). Neutrophils are pro-inflammatory cells, and the reduction of neutrophils has been reported to be key to breaking the inflammatory cycle in non-healing wounds [17].

**Figure 3:**
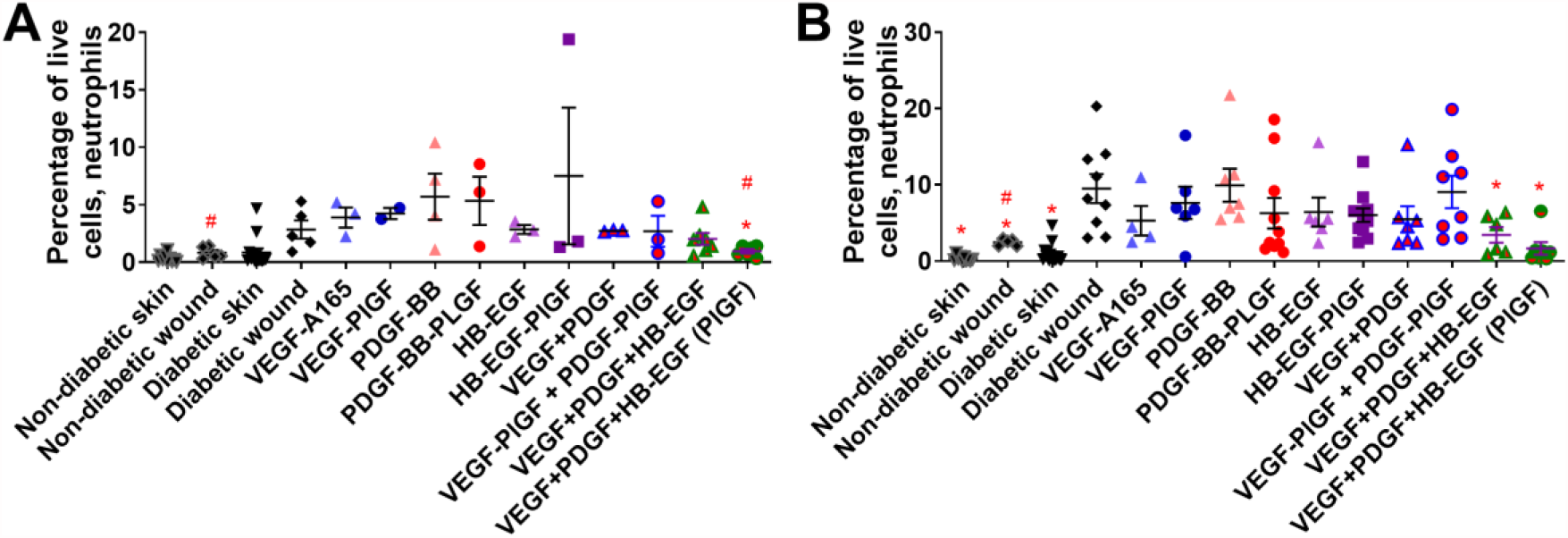
Neutrophil (CD45+, CD11b+, CD11c-, Ly6G+) count in wounds after 3-7 days of healing. (A) 3 days, (B) 7 days. Comparison includes unwounded skin from both the NOD and NOR mice. n ranges from 3 to 20. * denotes comparison to the diabetic wound. # denotes a comparison between untreated NOD wound and untreated NOR wound, and the WT GF(s) are compared to their counterpart Placental growth factor (-PLGF-2_123-144_) variant(s), e.g., VEGF vs VEGF-PlGF-2_123-144_. * = P < 0.05, P < 0.01, P < 0.001, Kruskal-Wallis + Mann-Whitney post-hoc test. # = P < 0.05, P < 0.01, P < 0.001, Mann-Whitney.

Following neutrophils (which can arrive at the wound in a matter of hours) are macrophages, which arrive and differentiate at the wound site over multiple days. Macrophages consist of many subtypes, some of which are inflammatory and others that are anti-inflammatory [17]. Treatment with the triple therapy increases the ratio of these anti-inflammatory macrophages to pro-inflammatory macrophages (Figure 4A and B). The -PlGF-2_123-144_ variant (VEGF-PlGF-2_123-144_, PDGF-BB-PlGF-2_123-144_, and HB-EGF-PlGF-2_123-144_) of the triple therapy increased the ratio compared to -WT variants (VEGF, PDGF-BB, and HB-EGF) of the triple therapy (Figure 4A and B).

**Figure 4:**
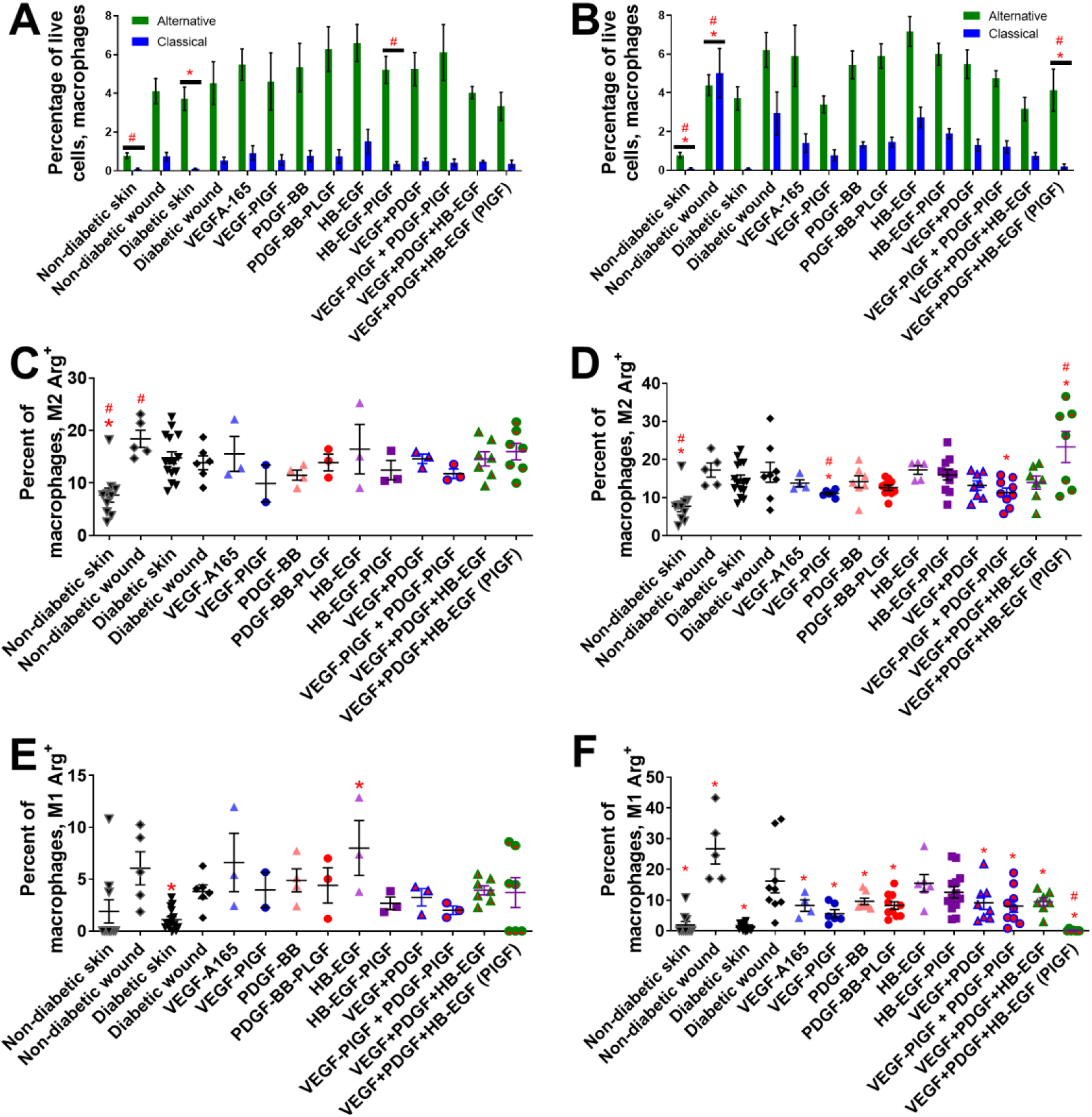
Macrophage cell count in wounds after 3-7 days of healing. Inflammatory M1 (CD45+, CD11b+, CD11c-, Ly6G-, Ly6C^low^, SSC^high^, MHC class II+) vs alternatively activated M2 (CD45+, CD11b+, CD11c-, Ly6G-, Ly6C^high^, SSC^low^, CD206+) macrophages after (A) 3 days or (B) 7 days. Arginase+ alternatively activated macrophages after (C) 3 days or (D) 7 days. Arginase+ classically activated macrophages after (E) 3 days or (F) 7 days. Comparison includes unwounded skin from both the NOD and NOR mice. n ranges from 3 to 20. * denotes comparison to the diabetic wound. # denotes a comparison between untreated NOD wound and untreated NOR wound, and the WT GF(s) are compared to their counterpart -PlGF-2_123-144_ variant(s), e.g., VEGF vs VEGF-PlGF-2_123-144_. * = P < 0.05, P < 0.01, P < 0.001, Kruskal-Wallis + Mann-Whitney post-hoc test. # = P < 0.05, P < 0.01, P < 0.001, Mann-Whitney.

Arginase is an enzymatic component of the urea cycle that is upregulated in wound-healing macrophages. Treatment with the -PlGF-2 variant triple therapy (VEGF-PlGF-2_123-144_, PDGF-BB-PlGF-2_123-144_, and EGF-PlGF-2_123-144_) increases the number of arginase positive wound healing macrophages at 7 days post wound healing (Figure 4D), as compared to diabetic wounds. The triple therapy (VEGF-PlGF-2_123-144_, PDGF-BB-PlGF-2_123-144_, and EGF-PlGF-2_123-144_) does not increase the number of arginase positive inflammatory M1 macrophages, and in fact significantly decreases the overall number of inflammatory M1 macrophages in wounds (Figure 4A and B).

Non-hematopoietic stromal cells are also key in wound healing. The triple therapy (VEGF-PlGF-2_123-144_, PDGF-BB-PlGF-2_123-144_, and EGF-PlGF-2_123-144_) did not increase the number of MSC after 3 days (Figure 5A), but increased the number of MSC at 7 days (Figure 5B) and increased the number of endothelial cells at both 3 and 7 days (Figure 5C and D). The -PlGF-2_123-144_ variant of triple therapy significantly increases the number of MSC and endothelial cells at 7 days compared to the WT variants (Figure 5B and D).

**Figure 5:**
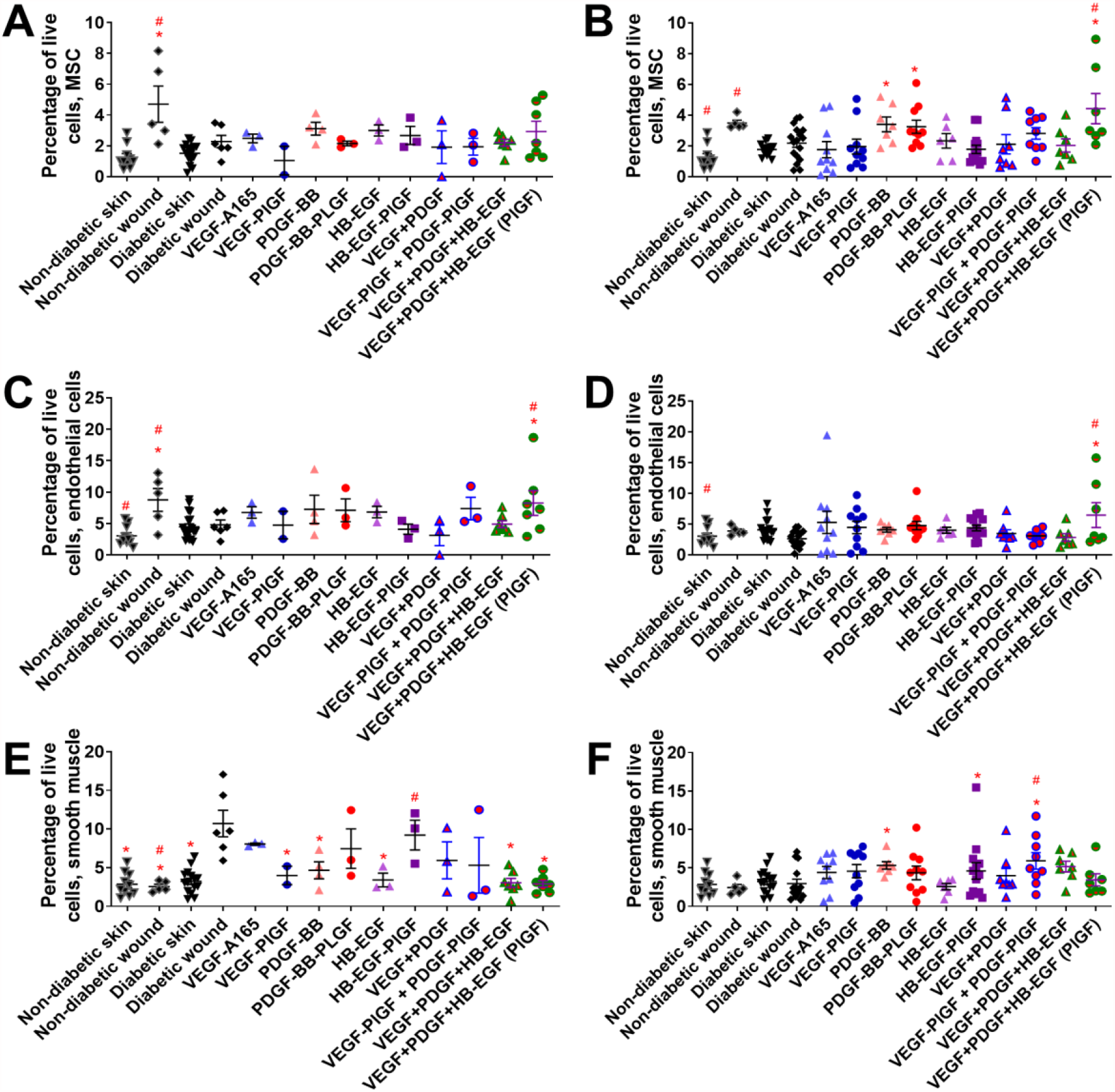
Non-hematopoietic cell count in wounds after 3-7 days of healing. Mesenchymal stem cells (CD45-, CD44+, CD29+, CD90+, SCA-1+) after (A) 3 days or (B) 7 days. Endothelial cells (CD45-, CD31+) after (C) 3 days or (D) 7 days. Smooth muscle cells CD45-, SMA+) after (E) 3 days or (F) 7 days. Comparison includes unwounded skin from both the NOD and NOR mice. n ranges from 3 to 20. * denotes comparison to the diabetic wound. # denotes a comparison between untreated NOD wound and untreated NOR wound, and the WT GF(s) are compared to their counterpart -PLGF-2_123-144_ variant(s), e.g., VEGF vs VEGF-PlGF-2_123-144_. * = P < 0.05, P < 0.01, P < 0.001, Kruskal-Wallis + Mann-Whitney post-hoc test. # = P < 0.05, P < 0.01, P < 0.001, Mann-Whitney.

MSC localization can also be used to measure the amount of vascularization in the wound environment, since MSC surround newly developed vascular tissue [47]. Immunoflourescence for MSC indicates an increase of MSC staining in triple-therapy treated wounds, though this increase is not significant (Figure S1). Treatment with triple therapy (VEGF-PlGF-2_123-144_, PDGF-BB-PlGF-2_123-144_, and EGF-PlGF-2_123-144_) also increased the number of proliferating cells in the wound significantly (Figure S2) when compared to both untreated diabetic wounds and to diabetic wounds treated with untargeted triple therapy.

The adaptive immune system participates in the later stages of wound healing [17]. Several growth factors (including the triple therapy) decrease the number of CD4^+^ T cells that are amphiregulin positive (Figure 6D). The triple therapy (VEGF, PDGF-BB, and HB-EGF) in both -WT and -PlGF-2_123-144_ variants decrease the number of effector T cells 1 week after treatment, with no significant difference between one another (Figure 7C). Treatment by GF or eGF do not significantly change the number of Tregs in the wound after 3 and 7 days (Figure 8A and B), though they do decrease the amounts of amphiregulin-positive cells after 3 and 7 days (Figure 8C and D).

**Figure 6:**
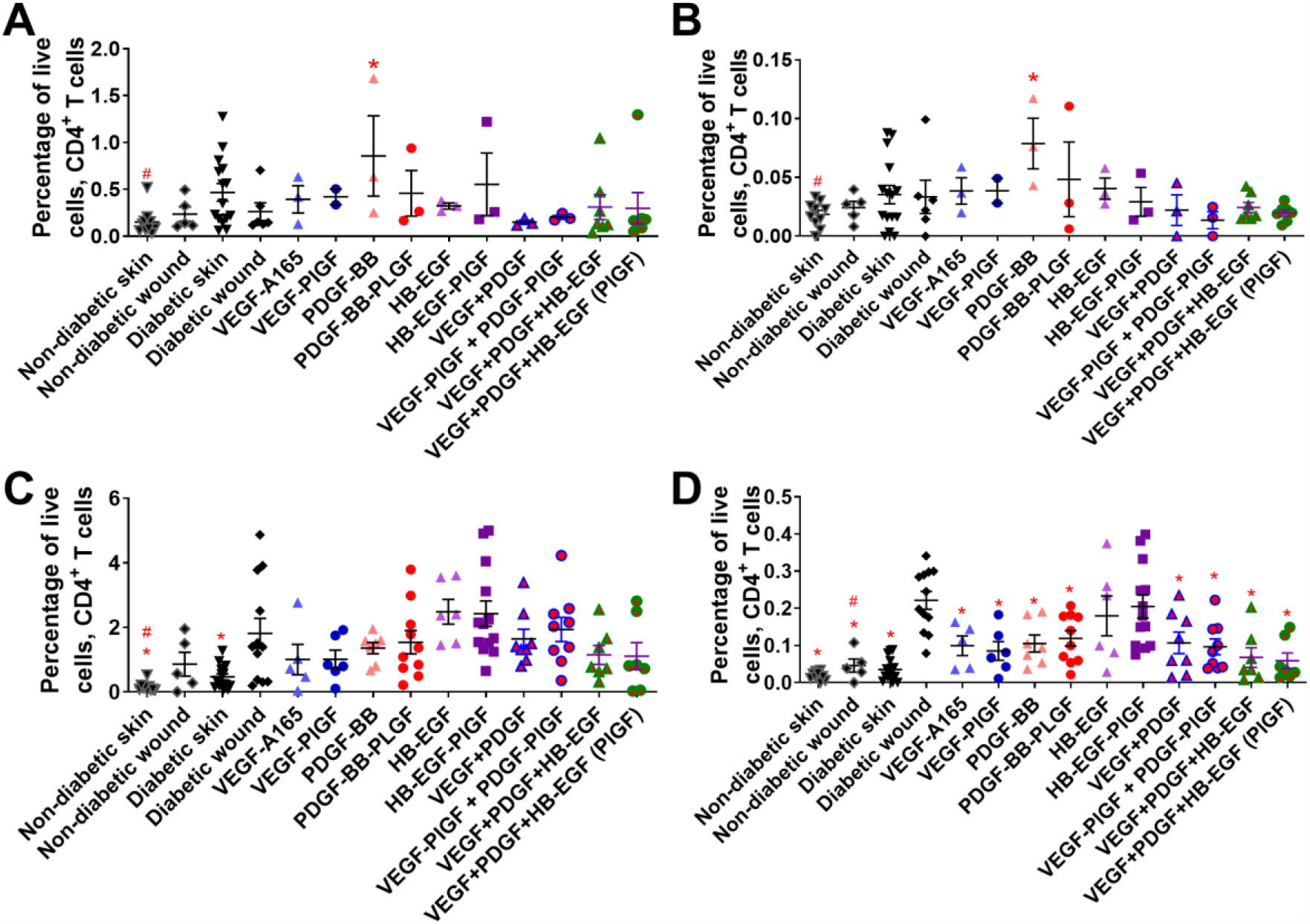
CD4+ T cell count in wounds after 3-7 days of healing. CD4+ T cells (CD45+, CD3+, CD4+) at (A) 3 days or (C) 7 days. Subset of CD4+ T cells that are amphiregulin positive at (B) 3 days or (D) 7 days. Comparison includes unwounded skin from both the NOD and NOR mice. n ranges from 3 to 20. * denotes comparison to the diabetic wound. # denotes a comparison between untreated NOD wound and untreated NOR wound, and the WT GF(s) are compared to their counterpart -PLGF-2_123-144_ variant(s), e.g., VEGF vs VEGF-PlGF-2_123-144_. * = P < 0.05, P < 0.01, P < 0.001, Kruskal-Wallis + Mann-Whitney post-hoc test. # = P < 0.05, P < 0.01, P < 0.001, Mann-Whitney.

**Figure 7:**
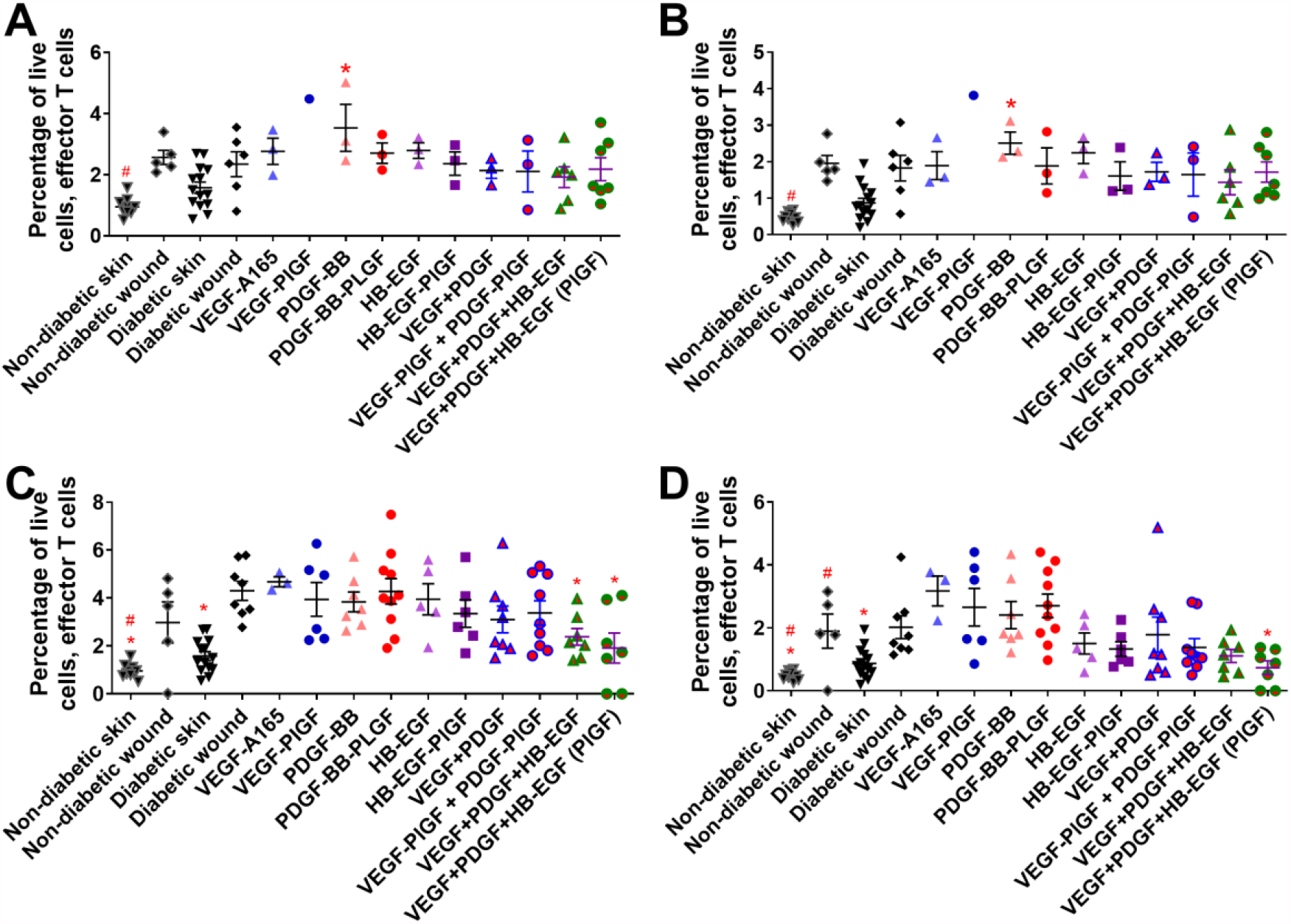
Effector T cell count in wounds after 3-7 days of healing. Effector T cells (CD45+, CD3+, CD44+, CD62L-) at (A) 3 days or (C) 7 days. Subset of effector T cells that are amphiregulin positive at (B) 3 days or (D) 7 days. Comparison includes unwounded skin from both the NOD and NOR mice. n ranges from 3 to 20. * denotes comparison to the diabetic wound. # denotes a comparison between untreated NOD wound and untreated NOR wound, and the WT GF(s) are compared to their counterpart -PLGF-2_123-144_ variant(s), e.g., VEGF vs VEGF-PlGF-2_123-144_. * = P < 0.05, P < 0.01, P < 0.001, ANOVA + Student t-test for post hoc. # = P < 0.05, P < 0.01, P < 0.001, Student’s t-test.

**Figure 8:**
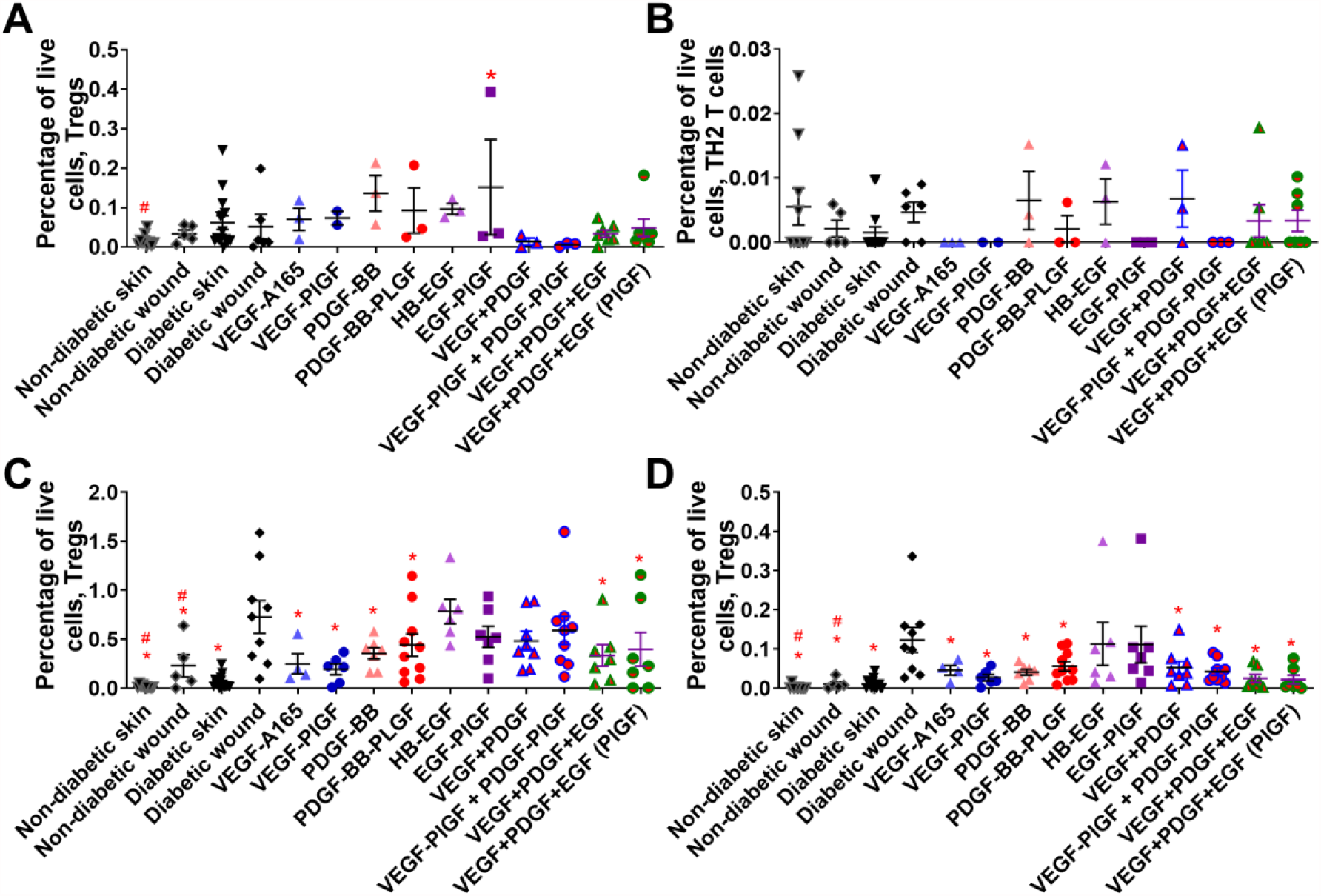
T regulatory cell count in wounds after 3-7 days of healing. T-regulatory cells (CD45+, CD3+, CD4+, CD25 high, Foxp3+) at (A) 3 days or (C) 7 days. Subset of effector T cells that are amphiregulin positive at (B) 3 days or (D) 7 days. Comparison includes unwounded skin from both the NOD and NOR mice. n ranges from 3 to 20. * denotes comparison to the diabetic wound. # denotes a comparison between untreated NOD wound and untreated NOR wound, and the WT GF(s) are compared to their counterpart -PLGF-2_123-144_ variant(s), e.g., VEGF vs VEGF-PlGF-2_123-144_. * = P < 0.05, P < 0.01, P < 0.001, Kruskal-Wallis + Mann-Whitney post-hoc test. # = P < 0.05, P < 0.01, P < 0.001, Mann-Whitney.

Since we treated the mice in a simulation of a walk-in clinic (where each mouse is treated as it individually becomes diabetic, and where each mouse is given insulin therapy adapted to its glycemic state), we can statistically compare the mouse-specific variables (age, previous glucose high level, glucose level during wound healing) and cell concentrations (neutrophils, macrophages, mesenchymal stem cells, epithelial cells, and effector T cells) with wound healing outcomes (granulation tissue and re-epithelialization) (Table 1). For a traditional linear regression model, the R^2^ value measures the percentage of variance in the response variable explained by the explanatory variable.

Interestingly, the variable with the highest R^2^ value for predicting wound healing outcome was the eGF treatment added. For wound extent the triple therapy treatment produced an adjusted R^2^ of 0.2279.

Individual cell types that significantly predicted wound healing outcomes were M2 macrophages and effector T cells, which both significantly correlated with increased granulation tissue (Table 1). Of the mouse-specific variables measured (mouse age, glucose average before wounding and before sacrifice, and whether the mouse achieved a glucose level of more than 450 mg/dl previously), none significantly predicted wound healing outcomes (wound closure or granulation tissue).

While the number of specific types of cells can correlate with wound healing outcomes in some cases, in other cases a combination of cell number and location is key to the wound healing outcomes. Multivariate linear regression models including both the treatment and the percentages of cells improved the predictive power for wound outcomes. For granulation tissue, the best model (with an R^2^ of 0.556, a substantial improvement over treatment alone which has an R^2^ of 0.17) used treatment, percentage of M2 macrophages, and percentage of neutrophils. Including interaction terms did not improve the R^2^ of the model. For wound extent, the best model (with an R^2^ of 0.32, a substantial improvement over treatment alone which has an R^2^ of 0.23) used treatment, percentage of effector T cells, and percentage of M1 macrophages. For wound extent, there is a significant interaction. Leaving out the interaction terms resulted in an R^2^ of only 0.26.

## Discussion

We previously demonstrated that co-administration of VEGF-A and PDGF-BB (and especially their -PlGF-2_123-144_ eGF variants) improves re-epithelialization in the mouse T2D db/db mouse model [20]. The present study considers three questions: first, whether the eGF triple therapy treatment (VEGF-A-PlGF-2_123-144_, PDGF-BB-PlGF-2_123-144_, and HB-EGF-PlGF-2_123-144_) has a statistically significant effect on wound healing for mice with T1D, second, whether addition of HB-EGF-PlGF-2_123-144_ to the dual combination was beneficial, and third, the specific way in which the cellular milieu of a wound affects healing upon treatment.

Here we demonstrate that eGF triple therapy significantly improves measurements both re-epithelialization and granulation tissue in the NOD mouse model of T1D, and that addition of HB-EGF-PlGF-2_123-144_ to the regimen is indeed beneficial (Figure 1). The eGF triple therapy also operates at doses far lower than previously approved GF therapeutics. Regranex’s dose of PDGF-BB for the treatment of diabetic foot ulcers is 7 µg/cm^2^ (or 1800 ng/28 mm^2^), administered daily for 2 weeks [36], is far greater than our dose of 200 ng/28 mm^2^, administered once.

The NOD model develops T1D via a similar immunological sequence as in human disease [48]. Our NOD colony was allowed to develop T1D without induction, permitting us to operate a model “walk-in wound clinic”, which resulted in data on the age, weight, and glucose history of each mouse in our study. These mouse-specific co-variates had less influence on the outcome of the wounds (re-epithelialization and granulation tissue) than the GF or eGF treatments (Table 1), or the cellular milieu of the wounds (Table 1), based on their R^2^ values. While mice age affects the hair growth of mice [49], we did not find a statistical relationship between mouse age and response to wounding (Table 1), suggesting the most important variables in wound healing were the growth factor treatment added to the wound, not the severity of the glucose dysregulation, the duration of the diabetes, or the age of the mice.

While healing wounds undergo changes in their cellular milieu [17], the cellular composition of a wound changes based on treatment with GFs and eGFs (Figures 3-8). This cellular milieu can be correlated with wound healing outcomes.

The cell populations with the highest R^2^ values for wound outcomes (re-epithelialization and granulation tissue) are M1 macrophages, M2 macrophages, and effector T cells (Table 1). Interestingly, while GF and eGF triple therapy treatment of wounds significantly affected the number of neutrophils and MSCs, neither of these cell types are predictive of improved wound outcomes on their own (Table 1). These findings contradict previous wound studies that have found MSCs to be integral to wound healing [50]. This discrepancy could be explained by the fact that several studies have found that MSCs are less functional is diabetic wounds, and that in these cases, additions of MSC-conditioned media may contribute to better wound outcomes [50].

eGF treatment significantly changed the ratio between M1 and M2 macrophages, but this change is not predictive of improved wound outcome (Table 1). These results could indicate that changes in the number of certain cells in a wound might be an outcome of improved wound healing, as opposed to a cause of improved wound healing.

Intriguingly, the opposite can also be true. The numbers of Tregs were not significantly increased by GF or eGF treatment, but Tregs did have a small but significant correlation with increased granulation tissue formation (Table 1). This indicates that triple therapy treatment improves wound healing using only some of the total constellation of available cell types and interactions within wounds. Depletion of Tregs slows wound closure in a non-diabetic model [46]. In the NOD diabetic mouse strain, dendritic cells are defective in their presentation to t cells, which could potentially explain why Tregs were not predictive of wound closure in our model [51]. Interestingly, dendritic cell function is restored in NOR mice [52].

Also interestingly, all 7 wounds treated with the triple therapy eGFs had zero M1 macrophages or zero effector T cells, suggesting that the lowered number of inflammatory cells might be key to improved wound outcomes. However, when correlated (Table 1), lower numbers of effector T cells correlate with improved re-epithelialization, and do not appear to affect granulation tissue formation. Lower numbers of M1 macrophages, conversely, correlate with increased granulation tissue formation, and do not appear to effect re-epithelialization.

Combining cellular data and treatment data (as variables) improved predictive outcomes for wound healing (Table 1). Since many factors go into successful wound healing, it is unreasonable to expect that any combination of explanatory variables would produce an R^2^ value of 1, i.e., completely explain the response variable. For multivariate regression models, additional explanatory variables can only ever increase the R^2^ value. To avoid models with too many explanatory variables, the adjusted R^2^ value includes a penalty term that increases with the number of variables. Hence, adding a useless variable will result in a smaller adjusted R^2^. Further, cross-validation ensured these models were not overfit to the data.

That the engineered growth factors (eGFs) with the -PlGF-2_123-144_ domain improve wound healing in comparison to the WT GFs at 7 days, but less so at 3 days, is mirrored in several of the datasets. The eGFs remain in the wound at higher concentrations at 1 week in comparison to the WT GF, and there is no significant difference at 3 days. There are also more significant differences in an eGF treated wound’s cellular milieu after 7 days of healing in comparison to the GF treated wounds, or NOD or NOR control wounds. The few differences in the wound’s cellular milieu at day 3 support the hypothesis that the longer duration of the eGFs in wounds improves wound healing, and that the individual eGFs are not intrinsically more potent signaling proteins that GF. These findings suggest that different cell types are responsible for different kinds of healing, either inducing re-epithelialization or increasing the granulation tissue in a wound.

While HB-EGF promotes wound re-epithelization [23], HB-EGF also effects keratinocyte migration and proliferation [30], depending on dose. Doses of 10-100 ng/ml slow keratinocyte proliferation [32], while the dose in this study is 200 ng per wound. Interestingly, immobilized HB-EGF preferentially causes keratinocyte migration, while soluble HB-EGF preferentially causes keratinocyte proliferation [31]. Since -PlGF causes ECM sequestration of growth factor, it’s possible that HB-EGF-PlGF-2_123-144_’s wound healing improvements may be related to the changes in the manner of HB-EGF-PlGF-2_123-144_’s presentation, in addition to changes in local dose concentration.

## Materials and Methods

### Fusion growth factor design

Wild-type human GFs (VEGF-A165, PDGF-BB, and HB-EGF) were selected for their varied effects on the wound environment, and their potential synergy for combination. Sequences obtained from Genscript (Piscataway, New Jersey) were transferred into pXLG [20], and the sequence for PLGF-2’s heparin binding domain (-PLGF-2_123-144_) was substituted for the native heparin-binding domains of VEGF-A165 and HB-EGF, and was added to the C-terminus of PDGF-BB, resulting in the growth factors denominated as VEGF-A-PlGF-2_123-144_, PDGF-BB-PlGF-2_123-144_, and HB-EGF-PlGF-2_123-144_. All growth factors were his-tagged at the N-terminus.

### Growth factor (GF) production

Wild-type and fusion GF were produced by transfection of human embryonic kidney HEK-293f cells, and purified by his tag affinity to a nickel column (Histrap, GE healthcare, Chicago, Illinois) via fast protein liquid chromatography (FPLC), as previously described [20]. GFs were assessed for purity by gel electrophoresis (>95%), and tested negative for endotoxins by a HEK-mouse TLR4 blue assay, as previously described [20]. GF concentration was assessed by bicinchoninic acid assay (BCA), according to the manufacturer’s directions (Thermo, Waltham, Massachusetts).

### Growth factor-receptor phosphorylation assay

Human endothelial cells from umbilical vein (ATCC, Manassas, Virginia) and MRC5 fibroblasts (ATCC) were serum starved overnight and stimulated with GFs, as previously described [20]. Cells were then lysed, and that lysate was assayed for GF receptor phosphorylation via DuoSet ELISA kits (R and D Systems, Minneapolis, Minnesota) for VEGFR2, PDGFR, and EGFR, as per the manufacturer’s instructions.

### Mouse walk-in clinic

The mice used in this study were non-obese diabetic (NOD) and control mice of the same background that do not develop diabetes (NOR). The mouse colony was maintained in-house, and blood glucose levels of the females were assessed weekly starting at 8 weeks old. Mice were defined as diabetic when they had two readings over 250 mg/dl or one reading over 400 mg/dl. To maintain glycemic control, albeit sub-optimally, diabetic mice were treated with 1-2 insulin pellets (Linshin, Scarborough, Ontario, Canada) per the manufacturer’s instructions. Specifically, the back of diabetic mice were shaved and sterilized with iodine and alcohol wipes. Following that, Linshin insulin pellets were placed in sharped 12 gauge needles and pellets were placed subcutaneously in the mouse back. One pellet was used for mice with glucose readings between 300 mg/dl and 500 mg/dl, and two pellets were used for mice with glucose readings greater than 500 mg/dl. In order to proceed to the wounding phase of the study, the mice had to have stabilized glucose levels between 250 mg/dl and 300 mg/dl for 2 subsequent weeks.

### Mouse wounding and healing

All animal experiments were performed with protocols approved by the University of Chicago IACUC. Wounding was performed as previously described [53]. Specifically, mice were anesthetized by 2% isoflurane. Mouse backs were shaved and sterilized with alcohol and iodine wipes. Two symmetrical wounds were made per mouse using a 6 mm biopsy punch. Sterile siliconized rubber splints were sewn onto the skin surrounding the wound using a suture kit. Wounds were then treated with GFs suspended in fibrin gel, bandaged with clear bandages, and wrapped with band aids to prevent scratching. Mice were monitored daily, and no signs of infection were observed during this study.

Wounds were resected after euthanasia (3 or 7 days of healing) with a 12 mm biopsy punch to keep the resection conditions consistent. Wounds were fixed in 4% paraformaldehyde, mounted in paraffin, and sectioned into 5 µm sections at the widest point of the wound. Wounds were then stained with hematoxylin and eosin. Wound extent and granulation tissue were assessed as previously described [20].

### Retention of GF in wounded tissue

Half of the full thickness wound was mechanically disrupted with a tissue homogenizer in the presence of protease inhibitors, as previously described [20]. Briefly, a 12 mm biopsy punch of a wound was cut in half with a razor blade. Half of the wound was frozen, and then wounds from the study were thawed, and mechanically ground in a tissue homogenizer (MP Biomedicals, Santa Ana, California) utilizing lysing matrix D (MP Biomedicals) in the presence of an EDTA-free protease inhibitor cocktail (Thermo). The amount of VEGF, PDGF-BB, and HB-EGF (WT and -PLGF-2_123-144_ variants) present in the tissue lysate were assessed by ELISA (DuoSet, R and D systems), per the manufacturer’s instructions.

### Flow cytometry

Half of a wound was mechanically disrupted by maceration, digested with 1 µg/ml collagenase-d (Sigma) for 1 hr with shaking, and frozen in 10% dimethylsulfoxide (DMSO). All samples were thawed and analyzed via flow cytometry at the same time, to create greater consistency within the flow cytometry data. Digested wound tissue was split into three equal portions and stained for flow cytometry based on panels for the following: non-hematopoietic cells, innate immune cells, and adaptive immune cells. Live dead stain was live-dead aqua, used per manufacturer’s instructions (ThermoFisher). Compensation was performed via UltraComp beads (ThermoFisher) per the manufacturer’s instructions. The panels used to identify cells were: neutrophils (CD45+, CD11b-, CD11c-Ly6G+), M1 inflammatory macrophages (CD45+, CD11b+, CD11c-, Ly6G-, Ly6C+, MHC class II+), M2 wound healing macrophages (CD45+, CD11b+, CD11c-, Ly6G-, Ly6C+, CD206+), mesenchymal stem cells (CD45-, CD44+, CD29+, CD90+, SCA-1+), endothelial cells (CD45+, CD31+), CD4 T cells (CD45+, CD3+, CD4+), CD8 T cells (CD45+, CD3+, CD8+), naï ve T cells (CD45+, CD3+, CD44-, CD62L+), central memory T cells (CD45+, CD3+, CD44+, CD62L+), effector T cells (CD45+, CD3+, CD44+, CD62L-), regulatory T cells (Treg; CD45+, CD3+, CD4+, CD25 high, Foxp3+), Th2: CD45+, CD3+, CD4, gata-3. Antibodies against arginase, amphiregulin and ki-67 were also added, where appropriate.

### Immunoflourescence

Paraffin-fixed 7-day wounds were sectioned to 5 mm. Antigen retrieval was performed using citrate buffer. Primary antibodies (CD73 (647), CD90 (488), [47]) were added at 5 mg/ml overnight. Wounds were washed 3x in PBS, and were exposed to DAPI (ThermoFisher) for 1 min. Cells were mounted using diamond anti-fade mounting media (ThermoFisher) to preserve fluorescence. Slides were imaged immediately using a confocal microscope (Olympus, Shinjuku City, Tokyo). All immunofluorescence wound sections were stained and imaged in the same batch, to better facilitate comparisons.

### Statistics

Wound healing was measured using two different response variables: the amount of granulation tissue (measured in mm^2^) and the wound re-epithelialization, i.e., the diameter at the widest point of the wound after 7 days of healing subtracted from the initial wound opening (measured in mm). As granulation tissue is related to the thickness of the wound, while wound extent is related to the wideness of the wound, these two response variables are not directly related to one-another. All statistics were run using the open-source computing program R.

To assess whether the triple eGF therapy treatment has a statistically significant effect on wound healing or the composition of cells in a wound (Figures 1, 3-8), we used ANOVA tests, or (for cases when the ANOVA conditions were not met), we used Kruskal-Wallis tests. This involved testing the normality of the wound outcome data (wound extent and granulation tissue), using treatment as the explanatory variable, and testing that groups had approximately equal variances. Our ANOVA tests compare the mean cell percentages for each treatment, while Kruskal-Wallis tests compare the mean ranks between groups. Next, we ran post hoc tests to compare each treatment to the control, using a two-sample t-test (if samples were approximately normally distributed) or a Mann-Whitney test (if normality was not satisfied). In all cases, Bonferroni corrections for multiple comparisons were used.

We also compared the outcomes of WT GF treatments to their corresponding eGF treatment (e.g., VEGF vs VEGF-PlGF-2_123-144_), denoted by # in Figures 1, 3-8. Each figure changes the response variable to a specific cell percentage. When the normality conditions were met, we carried out the comparison using two-sample t-tests (with independent populations). When the conditions were not met, we used Mann-Whitney tests.

To determine if the cellular milieu of a wound affects healing, we used exploratory data analysis, producing Table 1, followed by statistical tests to determine significance, with and without interaction terms. For each cell type (Figures 3-8), we fit linear models using the percentage of live cells as our explanatory variable and wound outcomes (wound extent or granulation tissue) as the response, using scatterplots to confirm the linear relationship. We then fit linear models with wound area as the response variable.

We included covariates including mouse age, average glucose levels at the time of surgery, and whether or not the mouse had >400 mg/dl glucose in their lifetime. For both response variables, we fit multivariate linear regression models (confirming linear relationships using scatterplots) with and without interaction terms, to investigate the significance of these covariates. We determined that none of the covariates were statistically significant on their own or in combination, that the effect size of the triple therapy treatment is far larger than that of the covariates, and that including interaction terms did not improve our models (except in cases of overfitting, identified and discarded using cross-validation), as measured by adjusted R^2^.

## Acknowledgements

We thank the Human Tissue Resource Center of the University of Chicago for histology analysis. We thank the Integrated Light Microscopy Core of the University of Chicago for Imaging.

## Abbreviations

(M2): Alternatively-activated macrophages
(M1): Classically-activated macrophages
(ECM): Extracellular matrix
(EGFR): Epidermal growth factor receptors
(GFs): Growth factors
(-PlGF-2_123-144_): Heparin-binding domain from placental growth factor-2
(HB-EGF): Heparin-binding epidermal-growth factor
(EGF-PlGF-2_123-144_): Heparin-binding epidermal-growth factor
(HER1): Human epidermal growth factor receptor
(PlGF-2_123-144_): Placental growth factor-2
(PDGF-BB): Platelet-derived growth factor-BB
(VEGF-PlGF-2_123-144_, PDGF-BB-PlGF-2_123-144_, and EGF-PlGF-2_123-144_): Triple therapy
(T1D): Type 1 diabetic
(T2D): Type 2 diabetes
(Tregs): T-regulatory cells
(VEGF-A): Vascular endothelial growth factor-A
(VEGFR2): VEGF-receptor-2
(WT): Wild-type

## Datasets

The datasets generated during the current study are available from the corresponding author on reasonable request

## Author’s contributions

MW, PSB, and JAH designed the project. MW performed experiments. MW and PB analyzed data. MW and JAH wrote the paper. All authors read and approved the final manuscript.

## Ethics approval

All the animal experiments performed in this work were approved by the Institutional Animal Care and Use Committee of the University of Chicago.

## Supplemental figures and legends

**Supplemental figure 1.**
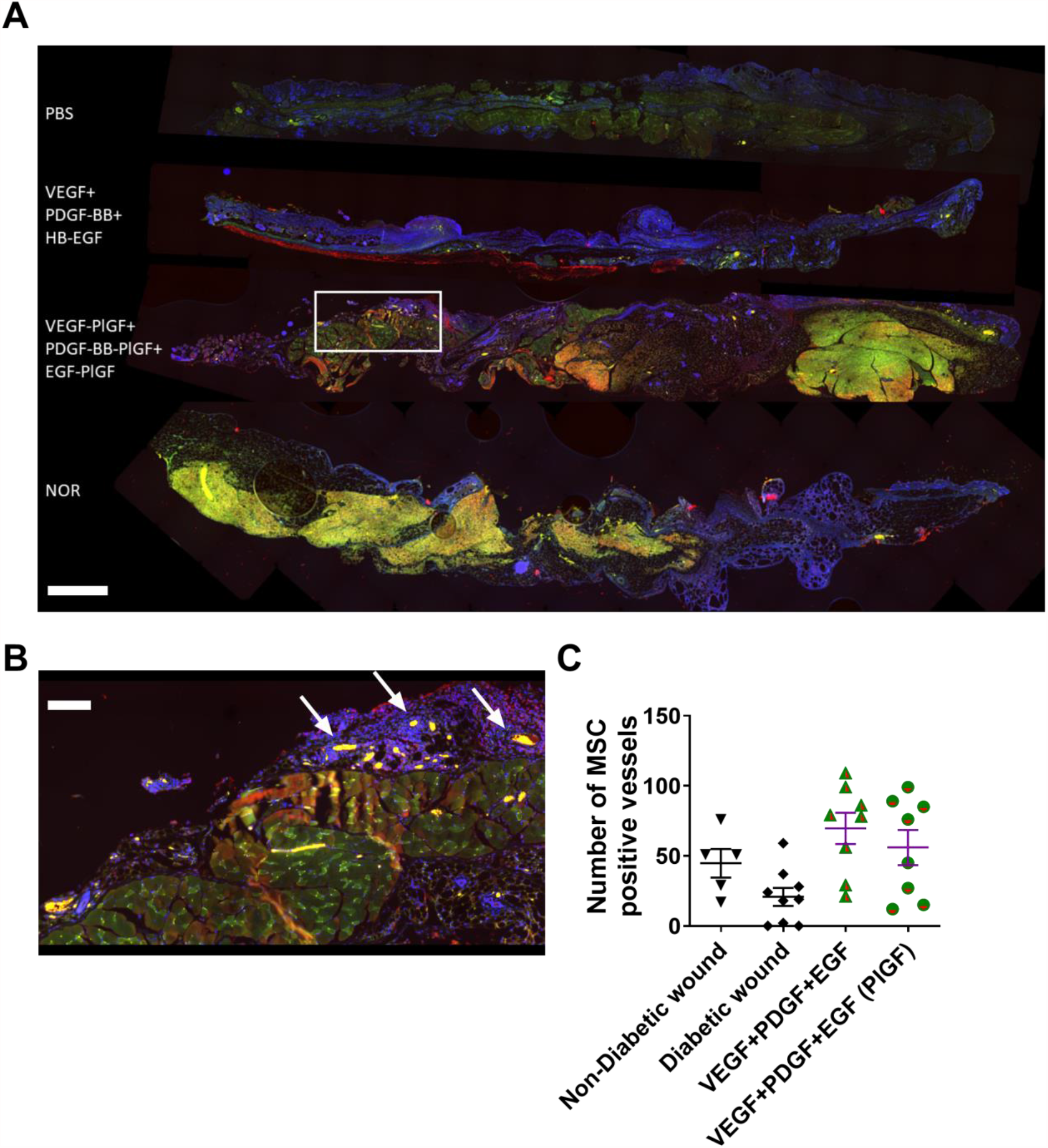
MSC staining in wounds. (A) Wounds with 7 days of healing were sectioned and stained for MSC using DAPI, CD90 (488), CD73 (647). (B) inset of A. Arrows indicate blood vessels for the purpose of counting, data in (C). There is no statistical significance in (C) by ANOVA or Student’s t-test. Red = CD73, green = CD90, blue = DAPI. Size bar = 1 mm, inset size bar = 100 µm.

**Supplemental Figure 2.**
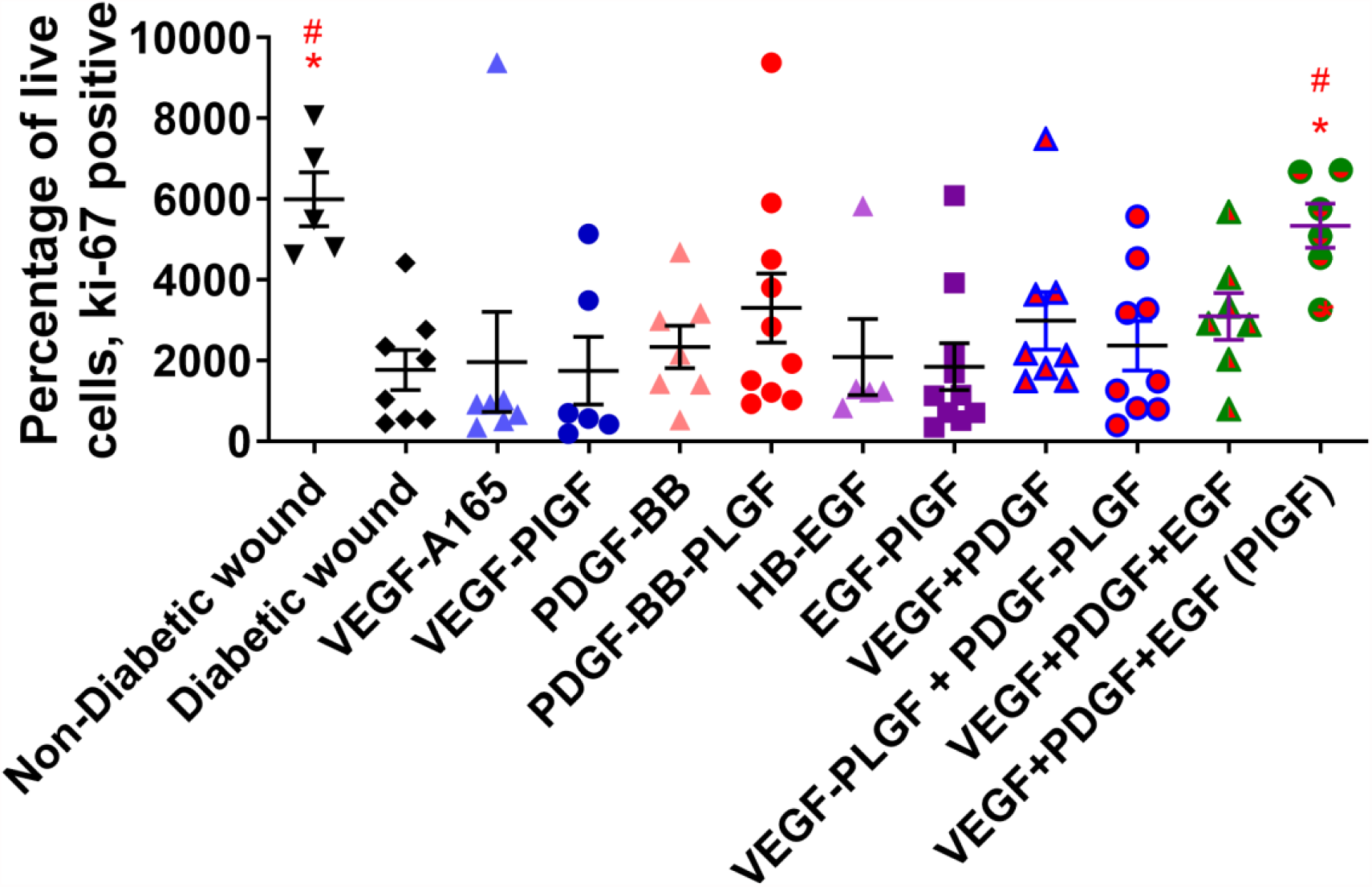
Treatment with triple therapy (VEGF-PlGF-2_123-144_, PDGF-BB-PlGF-2_123-144_, and EGF-PlGF-2_123-144_) increases the number of proliferating cells in wounds. Proliferating cells (ki-67) at 7 days healing. N ranges from 5 to 10. * denotes comparison to the diabetic wound. # denotes a comparison between untreated NOD wound and untreated NOR wound, and the WT GF(s) are compared to their counterpart -PLGF-2_123-144_ variant(s), e.g., VEGF vs VEGF-PlGF-2_123-144_. * = P < 0.05, P < 0.01, P < 0.001, 1 way ANOVA. # = P < 0.05, P < 0.01, P < 0.001, Student’s t-test,.

**Supplemental Figure 3.**
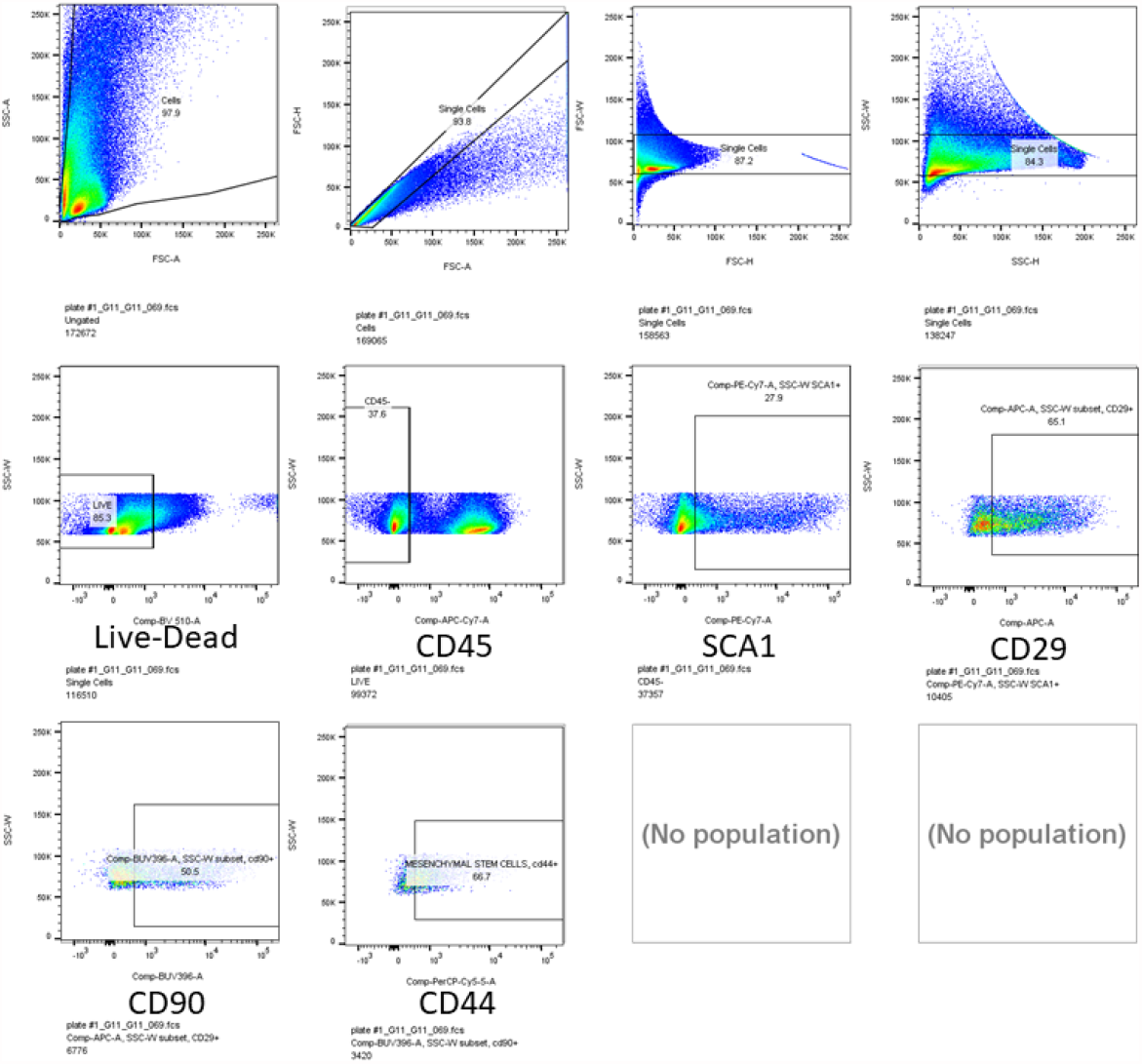
Gating strategy for MSC. Cells were gated for live cells using FSC-a vs SSC-a, followed by FCS-a vs FSC-h, followed by FSC-h vs FSC-w, followed by SSC-h vs SSC-w, then followed by live-dead aqua staining. We could expect 10,000+ cells per wound aliquot.

### Supplemental statistics note

This study was designed to answer two questions. First, whether the eGF triple therapy treatment (VEGF-A-PlGF-2_123-144_, PDGF-BB-PlGF-2_123-144_, and EGF-PlGF-2_123-144_) improves wound healing for mice with T1D in a statistically significant fashion, and second, whether changes in the cellular milieu of a wound affects healing.

We answered these two questions using two different response variables: the amount of granulation tissue (measured in mm^2^) and the wound extent, i.e., the diameter of the wound (measured in mm) at its widest point. As granulation tissue is related to the thickness of the wound, while wound extent is related to the wideness of the wound, these two response variables are not directly related to one-another.

Importantly, it is possible for an explanatory variable to be linearly related with both of them. We ran a large number of experiments, featuring different treatments (always including control mice) and measuring properties of the mice (age, etc.) as well as numbers of cells of different types. In each experiment, initial wounds were circular, with a diameter of 6 mm. Our experimental protocol followed the “walk-in clinic” model where mice would become diabetic spontaneously and only then be suitable to be included in an experiment. Consequently, different experiments had different numbers of mice, generally around 6 or 7 per experiment. Our statistics regarding the control group are aggregated from all experiments, as control mice always had the same experience. Hence, the control group contains 20 mice.

To answer whether the triple therapy treatment has a statistically significant effect on wound healing for mice with type I diabetes, we considered ten different treatment options, as well as a group of diabetic mice receiving no treatment, and a group of non-diabetic mice receiving no treatment. For each of the twelve groups, seven mice received the treatment and we measured the two response variables after seven days. We used an ANOVA test to compare these ten groups (Kruskal-Wallis for granulation tissue) and found that there was a statistically significant difference between them. We then conducted post-hoc tests (using Tukey’s HSD procedure) to compare each treatment to the untreated diabetic wound group.

As illustrated in Figure 1, several treatments were statistically significantly more effective than no treatment, with the largest impact being the triple therapy treatment. Furthermore, as expected, there was a statistically significant difference between the diabetic no-treatment group and the non-diabetic no-treatment group.

Next, we considered whether or not these differences were visible already on day three of the wound, rather than day seven. In this case, each group had n = 3 mice (so as to avoid giving an overly large number of mice treatments known to be suboptimal), though the triple therapy groups (-PlGF-2_123-144_ and WT) had n = 7 in each group, for reasons we will shortly explain. The result, shown in Figure 1, is that again the triple therapy was statistically significantly better than no treatment, for the response variable of wound extent (again using an ANOVA/Kruskal-Wallis test with appropriate post-hoc tests). We note that three days was not enough to see the difference for the response variable of wound area.

Next, we considered whether or not the addition of the -PlGF-2_123-144_ peptide to the GF (resulting in eGFs) was significant. Again, at both three days and seven days, we used Student’s t-tests (or Mann-Whitney tests) to compare the -PLGF-2_123-144_ variant of each treatment with the WT variants. As shown in Figure 1, at seven days, there is a statistically significant difference between the -PlGF-2_123-144_ eGF triple therapy treatment and the WT triple therapy treatment, for the response variable of wound extent. For wound area, and at three days, the different is not significant.

We turn now to the second question, regarding the cellular milieu. Using the wound sections from the previous experiments, with all twelve groups, at both three days and seven days, we measured the percentages of each of the types of cells discussed in the preceding sections. Figures 2 through 7 show these percentages, one figure per type of cell, for each of the twelve groups. As above, we carried out ANOVA/Kruskal-Wallis tests to determine if there were differences between the treatments, and appropriate post-hoc tests to compare each treatment to the diabetic non-treatment group. For each type of cell, we also compared eGF (ECM-super-affinity) and WT GF variants of each treatment. The results are discussed in the Results section.

Using R, we investigated whether percentages of each type of cell had predictive power for wound area and wound extent. For each cell type, we fit linear models using the percentage of live cells as our explanatory variable and wound extent as the response, again using scatterplots to confirm the linear relationship. We then fit linear models with wound area as the response variable. For each model, we recorded the adjusted R^2^ value and the p-value, and these are displayed in Table 1. For wound extent, the percentage of effector T cells was a statistically significant predictor, with an adjusted R^2^ of 0.036. For wound area, the percentage of M2 macrophages (and the percentage with arginase) were statistically significant predictors, with R^2^ of 0.08 (respectively, 0.058). We caution that the p-values in Table 1 should be interpreted with caution, as so many models were fit. However, we were cautious not to include models that were overfit to our data. We used cross-validation to detect and discard such models, including models with higher-order interaction terms, or too many explanatory variables. As the present paper is the first to consider the question of the predictive power of live cell percentages for wound healing, we hope subsequent papers will confirm the relationships we identify as significant in the current exploratory analysis.

